# Tandem duplication of serpin genes yields functional variation and snake venom inhibitors

**DOI:** 10.1101/2025.01.07.631777

**Authors:** Meilyn S. Ward, Matthew L. Holding, Laura M. Haynes, David Ginsburg

## Abstract

Tandem duplication of genes can play a critical role in the evolution of functional novelty, but our understanding is limited concerning gene duplication’s role in coevolution between species. Much is known about the evolution and function of tandemly duplicated snake venom genes, however the potential of gene duplication to fuel venom resistance within prey species is poorly understood. In this study, we characterize patterns of gene duplication of the SERPINA subfamily of genes across in vertebrates and experimentally characterize functional variation in the SERPINA3-like paralogs of a wild rodent. We find the hallmarks of rapid birth-death evolution of SERPINA1-like and SERPINA3-like genes within and between rodent lineages. Next, we recombinantly expressed the 2 paralogous duplicates of SERPINA1 and 12 paralogous duplicates of SERPINA3 found in the genome of the big-eared woodrat (*Neotoma macrotis*), a species known to be resistant to protease-rich rattlesnake venoms. We found that two SERPINA3 paralogs inhibit snake venom serine proteases, indicating that these proteins have potential as resistance factors in SERPIN-mediated venom resistance. In addition, functional variation is apparent among paralogs, including neofunctionalization to inhibit both chymotrypsin-like and and trypsin-like proteases simultaneously for one venom-inhibiting paralog. Our results provide further evidence that the rapid evolution of SERPINA1 and SERPINA3 gene copy number across rodents has adaptive potential by producing functionally-diverse inhibitors.

## Introduction

Coevolution governs many interactions in nature, from mutualism to parasitism and predator-prey relationships (Holding et al., 2016a; Thompson, 2005; Z. Wu et al., 2023). The escalating “arms race” is a common metaphor for the latter case especially, describing pairs of morphological, physiological, and molecular traits that evolve in response to changes in the other in coevolving species (Benkman et al., 2003; Brodie et al., 2002; Toju, 2008). Comparative genomics, aided by the emergence of high-quality contiguous genomes for many species, has uncovered the common role that tandem gene duplication plays in adaptive evolution to environmental challenges (Brovkina et al., 2023; Demuth and Hahn, 2009; Hahn, 2009; Lallemand et al., 2020). During tandem duplication, the number of copies of a gene can change the effective dose of that protein available for use or allow more leniency for non-synonymous mutations that can eventually lead to functional diversification of copies. Genome-wide, tandem duplications and resulting species-specific copy number variation are enriched for genes directly interacting with the outside world rather than developmental or homeostatic genes (Brovkina et al., 2023; Demuth and Hahn, 2009). Therefore, multi-component, protein-based molecular arms races between species may be fueled by tandem duplication, the characterization of which would enrich our understanding of how coevolution fuels genomic diversity and functional divergence of genes.

Animal venoms are prime examples of adaptation fueled by tandem duplications because of their essential role in the lives—and deaths—of both predator and prey. Snake venoms are among the most well-understood venoms and have several gene families that have undergone tandem duplication (Casewell et al., 2013; Dowell et al., 2016). Snake venom serine proteases (SVSPs) are one such protein family in which genes have been duplicated several times, allowing for great diversity within and between species. From an ancestral duplication of the glandular kallikrein (KLK1) gene, SVSPs have diversified into gene copies with specificity for diverse targets, yielding proteins with potent pharmacological effects and even translational value (Itoh et al., 1988). They have a wide array of functions, many of which affect the physiology of hemostasis in the envenomated species (Markland, 1998; Serrano, 2013).

Consistent with a coevolving system, many natural prey species are highly resistant to snake venom. Wild rodents, for example, are able to survive venom doses hundreds to thousands of times greater than that which would kill a lab mouse (*Mus musculus*) (Balchan et al., 2024; Perez et al., 1978). Despite the remarkable levels of resistance, we know very little about the molecular basis of resistance to the many venom proteases and how resistance and venom coevolve. Previously identified resistance factors <u>(reviewed in Holding et al., 2016)</u> largely include generic protease inhibitors such as alpha-2-macroglobulin and inter-alpha-trypsin inhibitor (Biardi et al., 2000; de Wit and Weström, 1987; Omori-Satoh et al., 2000).

A possible source of a more specific inhibitor of snake venom proteins is the serine protease inhibitor, or SERPIN, protein superfamily, which contains members that have previously been implicated in venom resistance (Barbour et al., 2002a; Gibbs et al., 2020). SERPINs share a fishing bait-like reactive center loop (RCL) that extends above the main globular body of the SERPIN and, in inhibitory SERPINs, mimics the target sequence of the protease(s) it inhibits (Gettins and Olson, 2009). Within the RCL, residues are traditionally labeled based on their location relative to the cleavage site, with the residue on the N-terminal side of the cleavage site being the P1 residue and that on the C-terminal side being the P1’ residue. When a protease attempts to cleave the RCL of a SERPIN, the SERPIN will attempt to capture and inhibit the protease by forming an acyl intermediate and inserting the RCL into a central β-sheet of the globular domain. Stabilization of the acyl intermediate results in permanent covalent attachment and permanent loss of function for both the protease and the SERPIN (Gettins, 2002; Huntington, 2011; Lawrence et al., 1995). Alternatively, if hydrolysis of the acyl intermediate occurs before insertion into the central β-sheet then the SERPIN is cleaved and rendered nonfunctional while the protease retains function.

Because of SERPINs’ 1:1 inhibitory mechanism and the target-specific mimicry of the RCL, gene duplications of SERPINs can either increase the dose of available inhibitors or allow for mutational flexibility to target orthogonal proteases. Gene duplications have been found within the vertebrate-specific SERPINA subfamily of SERPINs, including α-1-antitrypsin (SERPINA1) and α-1-antichymotrypsin (SERPINA3) in rodents (Barbour et al., 2002b; Borriello and Krauter, 1991; Heit et al., 2013). SERPINA1 has also been found to be evolving at a higher rate than the background in ground squirrels (Gibbs et al., 2020; Ochoa et al., 2023). However, SERPINA1 is not the only SERPINA to see such expansion in rodents. In the same tandem array, SERPINA3 is also duplicated in many rodents (Forsyth et al., 2003). Prior work with SERPINA1 paralogs in mice observed differential protease inhibition by each (Barbour et al., 2002a). One compelling theory for the function of SERPINA paralogs is that they serve as inhibitors of SVSPs in snake predators. Squirrel SERPINA1 was enriched by bait-capture of snake venom and was found to be enriched in tissue after envenomation (Cavalcante et al., 2022; Gibbs et al., 2020). In addition, paralogs of SERPINA1 from *Mus saxicola* were found to differentially form inhibitory complexes with snake venom proteases (Barbour et al., 2002a).

Here, we combine comparative genomics and direct functional tests to identify the lineages in which SERPINA duplications occur most frequently and then establish whether these duplications are producing functional variation in protease inhibition in non-model rodents. We specifically test the hypothesis that tandem duplication of SERPINA3s of *N. macrotis*, a wild rodent belonging to a genus known to have significant resistance to snake venom, is an adaptation to inhibit the serine proteases in the venom of its native rattlesnake predators (de Wit, 1982; Perez et al., 1978; Robinson et al., 2021).

## Methods

### Genome annotation

Chromosome-scale or large-scaffold-scale genomes were downloaded from NCBI’s Genome resource (Sayers et al., 2022). SERPINA protein sequences from UniProt (The UniProt Consortium, 2023) were queried with tblastn (Altschul et al., 1990) with e-value of 10^-6^ against each genome to locate putative exon hits for SERPINA genes. A total of 68 genomes were manually re-annotated at SERPINA tandem arrayed locus, including mammals, reptiles, birds, and fish. Annotated SERPINA coding sequences were translated and web blastp searches with default settings were used to establish homology to other annotated sequences for nomenclature in our study. For the putatively novel SERPINA genes in the genomes of the woodrats *Neotoma fuscipes* and *N. macrotis*, we further refined the coding sequence annotation using RNA-seq reads visualized with splice-aware alignment in HiSAT2 <u>(Kim et al., 2019)</u>(Kim et al., 2019) . We then named the paralogs of SERPINA1 and SERPINA3 according to their genomic order on the chromosome assembly (e.g. SERPINA3-1, SERPINA3-2, … SERPINA3-12). To assist the study of gene order relationships over evolutionary time, we applied similar nomenclature to our annotations of SERPINS in other species.

### Phylogenetic analyses

The coding sequences used to create the phylogeny were taken from the tandem array that, in humans, is on chromosome 14 between ppp4r4 and gsc. Other SERPINAs are present on other chromosomes in humans and many other animals, but we chose to focus on this tandem array’s evolution and function. To recover a species-tree aware phylogenetic reconstruction of the SERPINA gene tree, we first generated a codon-alignment by aligning the coding sequences using MUSCLE (Edgar, 2004) with kmer6_6 distance in the first iteration and pctid_kimura distance the subsequent iterations (200 max iterations), using the CLUSTALW weight scheme (Thompson et al., 1994), then back-translating. Next, we used timetree.org (Kumar et al., 2022) to obtain a dated species-tree for the 67 species present in our alignment. We then used GeneRax v2.1.3 (Morel et al., 2020) to produce a gene tree and reconcile it to the species tree with the reconciliation model set to ‘UndatedDL’ to infer duplication and loss events. Finally, we used ThirdKind v3.5.0 (Penel et al., 2022) to visualize the reconciled trees.

### Bacterial expression and purification

Each gene of interest was inserted into a pET-24(+) plasmid vector (Fig. S1), with a Strep tag on the N-terminal side, as well as a FLAG tag then a 6-His tag on the C-terminal side. Signal peptides were identified using signalP5 (Almagro Armenteros et al., 2019) and removed. In SERPINA3-5 and SERPINA3-6, no signal peptide was detected and thus no alterations were made to the coding sequence. Each tag was separated from other tags or the gene sequence by a Gly-Ser-Ser-Gly flexible linker. Constructs were synthesized at Twist Bioscience (South San Francisco, CA, USA). These vectors were chemically transformed into NiCo21(DE3) Competent *E. coli* (New England BioLabs Inc., Ipswich, MA, USA; #C2529H), selected with 30 ug/mL kanamycin.

*E. coli* were cultured in LB broth at 37℃ to an optical density of 0.6-0.9 abs at 600 nm, followed by 0.4 mM IPTG to induce expression before further incubation at 37℃ for 2 hours. Cells were recovered by centrifugation at 6000 x g for 20 minutes at 4℃, washed with 0.85% NaCl, pelleted again, and resuspended in 1 mL HBS (25 mM HEPES, 150 mM NaCl, pH 7.4) for every 100 mL of the culture. To prepare lysate, 70 uM lysozyme, 110nM DNaseI, 5 mM MgCl2, and 0.5 mM CaCl2 were added and incubated while rotating for 30 minutes at room temperature. Three freeze-thaw cycles were performed with liquid nitrogen, then the lysate was cleared by centrifuge at 16000 x g for 20 minutes at 4℃, passed through a 0.22 μm filter, and diluted to 1:4 in 5 mM imidazole (BioUltra, ≥99.5%; Sigma, Burlington, MA, USA, cat. # 56749-250G) in HBS.

Purification was completed using fast protein liquid chromatography (FPLC) on an AKTÄ Pure FPLC system (Cytiva Corp., MA, USA) with a HiTrap TALON crude 1 mL column (Cytiva Corp.). The column was equilibrated with HBS with 5 mM imidazole. Lysate was then applied and washed with 7.5 mM imidazole in HBS until the UV measurements were stable (within 0.1 mAU) for 1 minute. Elution was carried out with 150 mM imidazole in HBS. Buffer exchange into HBS was conducted on a HiPrep™ 26/10 Desalting column, and fractions containing the peak were pooled, aliquoted, and stored at -80L.

Purity was assessed on Invitrogen Novex Wedgewell™ 4-20% Tris-Glycine polyacrylamide gels (cat# XP04205), loading 1.2 ug of total protein into each well and staining with Invitrogen SimplyBlue SafeStain. To differentiate target proteins from low abundance impurities, a western blot was performed, using the Coomassie-stained gels. A three-step dry transfer process was used, beginning at 20 V for 1.5 minutes, then 23 V for 6 minutes, and finally 25 V for 3 minutes. The membranes were blocked with 2% bovine serum albumin (BSA) (Sigma-Aldrich, cat# A7906) in TBS-T (20 mM Tris, 500 mM NaCl, pH 7.5; with 0.1% Tween-20). Western blots were probed with a primary mouse anti-Strep-II tag antibody (Abcam, Inc; Cambridge, UK, AB184224, Lot #1033268-2) at a 1:1000 dilution in TBS-T for two hours, followed by a secondary goat anti-mouse HRP-conjugated antibody (1:5000 dilution) for one hour (BioRad, cat# 170-6516). Three fifteen-minute washes in TBS-T were conducted between antibody applications and before addition of the chemiluminescent substrate (SuperSignal™ West Femto Maximum Sensitivity Substrate, Thermo Scientific, #34095). The chemiluminescent substrate was applied for 3 minutes before imaging by a ChemiDoc™ MP (BioRad) at auto-optimal exposure. For the A3 paralogs, 1 ug was loaded and for the A1 paralogs, 1.2 ug was loaded.

### Enrichment and biotinylation of snake venom serine proteases

Lyophilized whole venom was diluted to 20 mg/mL in HBS and applied to a HiTrap Benzamidine FF(HS; Cytiva Corp.) columns equilibrated with 0.05 M Tris-HCl containing 0.5 M NaCl, pH 7.4. Fractions of 0.5 mLs were eluted with 0.05 M glycine, pH 3.0 and neutralized with 7.5uL 1M Tris-HCl, pH9. Final pH of fractions was approximately 7.4. Fractions with concentration of >0.150 mg/mL were pooled then buffer-exchanged into 25 mM HEPES containing 150 mM NaCl, pH 7.4.

Enriched SVSP fractions were biotinylated for sensitive detection in western blots of the serine protease mixture. Biotin was attached to SVSPs using the EZ-Link Micro Sulfo-NHS-Biotinylation Kit (Thermo Scientific, MA, USA; Cat# 21925) with a 20 mM fold-excess of biotin and approximately 1mg of input venom protein. Biotinylated venom was buffer exchanged and recovered using a 5mL Zeba Spin Desalting Column (Thermo Scientific; Cat. #89891) and stored at -80C until use. Detection of biotinylated venoms was achieved through western blot as described above, but using a single step incubation with HRP-conjugated streptavidin diluted 1:7500 in TBS-T.

### Reactions of SERPINs and proteases

Functional variation was assessed by visualizing inhibition of the proteases cathepsin G, chymotrypsin, trypsin, and elastase, the human substrates of human SERPINA1 and SERPINA3. These proteases are the known targets of human Serpina1 and Serpina3, and are here used as a reporter in the absence of identified rodent-specific targets. Proteases and SERPINs were mixed at a 2:5 molar ratio, respectively (0.9 μM protease and 2.25 μM SERPIN). Proteins were incubated at 22L on an Eppendorf Thermomixer R at 300 rpm. Empirically determined incubation times were as follows: SERPIN-chymotrypsin mixtures were incubated for 2 minutes, the SERPIN-trypsin mixtures were incubated for 1 minute, the SERPIN-elastase mixtures were incubated for 5 minutes, and the SERPIN-cathepsin G mixtures were incubated for 5 minutes. Mixtures were quenched with 4X Laemmli Blue buffer (BioRad, Cat. #1610747) containing 10% 2-Beta-mercaptoethanol as a reducing agent and immediately boiled at 95L for 10 minutes before SDS-PAGE.

For mixtures of SERPINs and benzamidine-purified SVSPs, venom proteases and SERPINA3 paralogs were again mixed at a 2:5 ratio), respectively, and incubated at 37L for 20 minutes. As before, reactions were quenched and boiled. Gel samples were run on Invitrogen Novex™ Wedgewell™ 10% Tris-Glycine Plus polyacrylamide gels (cat# XP00105). The biotinylated venom and Strep-II tag labeled SERPIN were visualized simultaneously with a western blot. The membranes were blocked in 2% BSA in TBS-T (0.01% Tween-20) overnight, then probed with 1:7500 N100, HRP-conjugated streptavidin and developed with chemiluminescent substrate (SuperSignal™ West Femto Maximum Sensitivity Substrate). To optimize the imaging process for detecting the complexes of interest in the presence of greater amounts of unbound protein, the portion of each blot above the 50kDa marker was imaged separately at a longer exposure.

In addition to the benzamidine-purified SVSPs, the activity of SERPINA paralogs was tested against two commercially purified SVSPs, Russell’s viper venom factor V activator (RVV-V) (Haematologic Technologies, Inc., Essex Junction, VT, USA, cat# RVVV-2000) and *A. contortrix* Protein C Activator (Protac) (Enzyme Research Laboratories, South Bend, IN, USA, cat# 113-01). SERPINA paralogs were mixed with these SVSPs at a 1:1 molar ratio and incubated at 37L for 20 min then quenched, boiled, run on 4-20% Tris-Glycine polyacrylamide gels, stained with Invitrogen SimplyBlue™ SafeStain, and imaged.

## Results

The tree containing all SERPINA paralogs within the tandem array (Fig. 1) displays the pattern of expansion events that have occurred within the SERPINA family in rodents. Fish and non-mammal tetrapods have SERPINA1-like, A3-like, and A10-like genes, whereas mammals were unique in the presence of many single-copy SERPINA subfamily genes. Gene duplications are most frequent in SERPINA1-like and SERPINA3-like gene clusters in mammals (Supp. Table 1). Clades composed wholly of SERPINA1 and SERPINA10 are present in every class of animal represented in this tree, suggesting these formed from duplication early in vertebrate evolution and copies have been maintained in daughter lineages.

**Figure 1.**
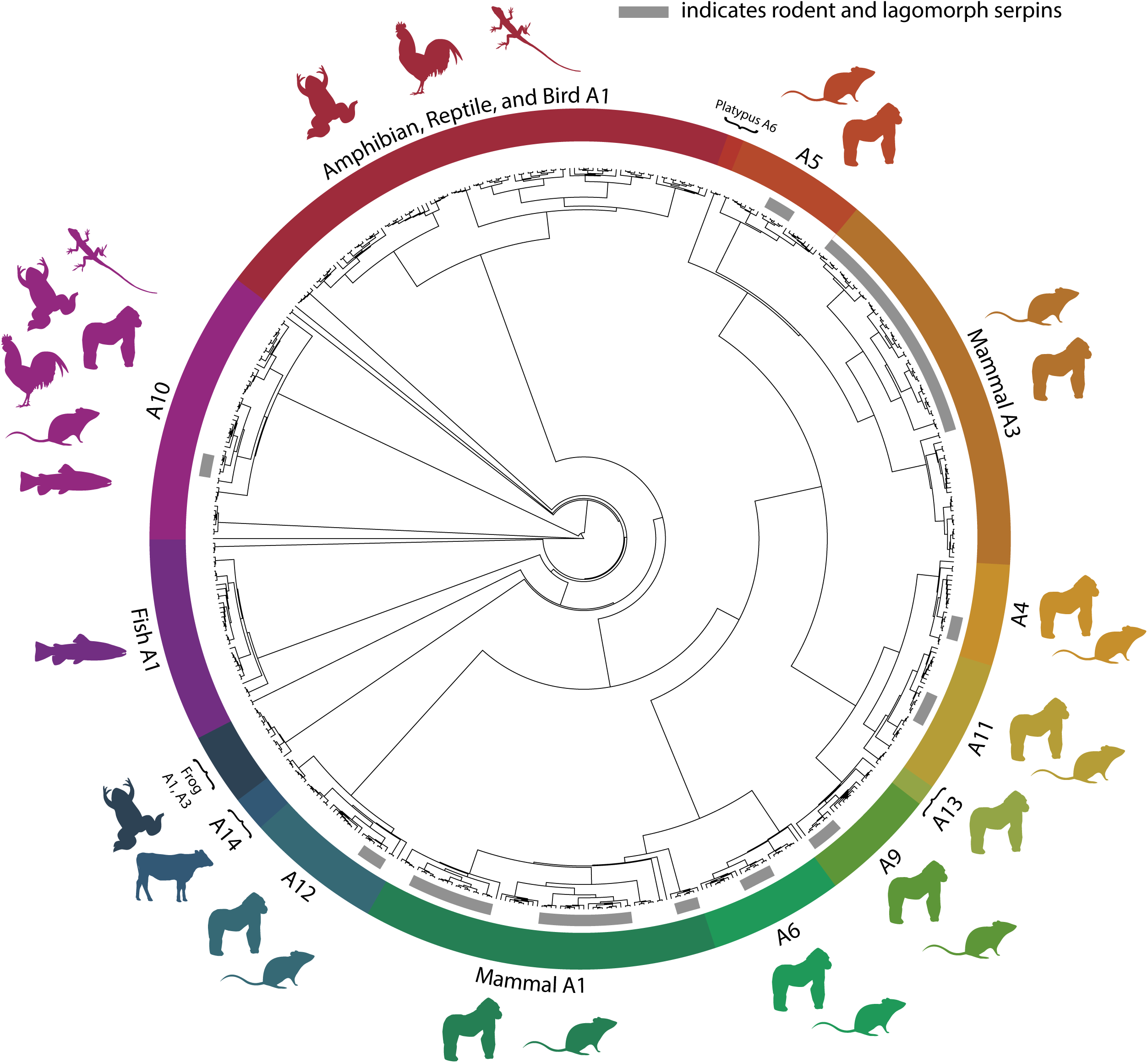
Evolution of A-clade SERPIN genes at the tandemly-arrayed locus. Shown is a maximum-likelihood phylogenetic tree of members of the SERPINA tandem array bounded by ppp4r4 and gsc genes. The colored segments of the circle represent different SERPINs, and the animal silhouettes adjacent to each segment show in which genome(s) that gene is present (amphibian, reptile, fish, bird, non-primate mammal, and primate). The grey inner arcs represent rodent and lagomorph paralogs of each gene, with the length of each arc being proportional to the number of duplications; there has been significant duplication of SERPINA1 and SERPINA3 compared to other animals, as well as other genes in the same tandem array. SERPINA10 was present in every class of animal represented in this study. Of note, SERPINA13 is unique to non-human apes and monkeys. Likewise, SERPINA14 is unique to non-rodent, non-simian mammals.

From the reconciled gene tree and species tree (Fig. 2a, 2b), patterns of birth-death evolution in the gene family can be inferred. Rodents and related lagomorph species stand out in terms of copy number relative to other clades. The two species within the *Neotoma* genus share most of their duplication events, likely reflecting their recent divergence. In addition, the expansions occur differently across rodent genera. *Mus musculus* experienced four duplications each of SERPINA1 and SERPINA3, while *Rattus norvegicus* only had two duplications of SERPINA3. The four ancestral duplication events uniquely shared by these two species all occurred within SERPINA3s. New World rodents experienced three shared ancestral duplication events, all of which are sequential duplications stemming from SERPINA3. In *Neotoma*, each of these duplications are repeated several more times, eventually resulting in the twelve total copies of SERPINA3 present. The most recent of the New World duplication events produced three of the *N. macrotis* SERPINA3 paralogs found to successfully inhibit proteases (further discussed later). Quantitatively, gene duplications and losses were on average higher in rodents and related lagomorphs than in other vertebrate species (p = 0.068; p = 0.041, respectively; Fig. 2c, 2d), indicating an overall elevated role of birth-death evolution in these lineages.

**Figure 2.**
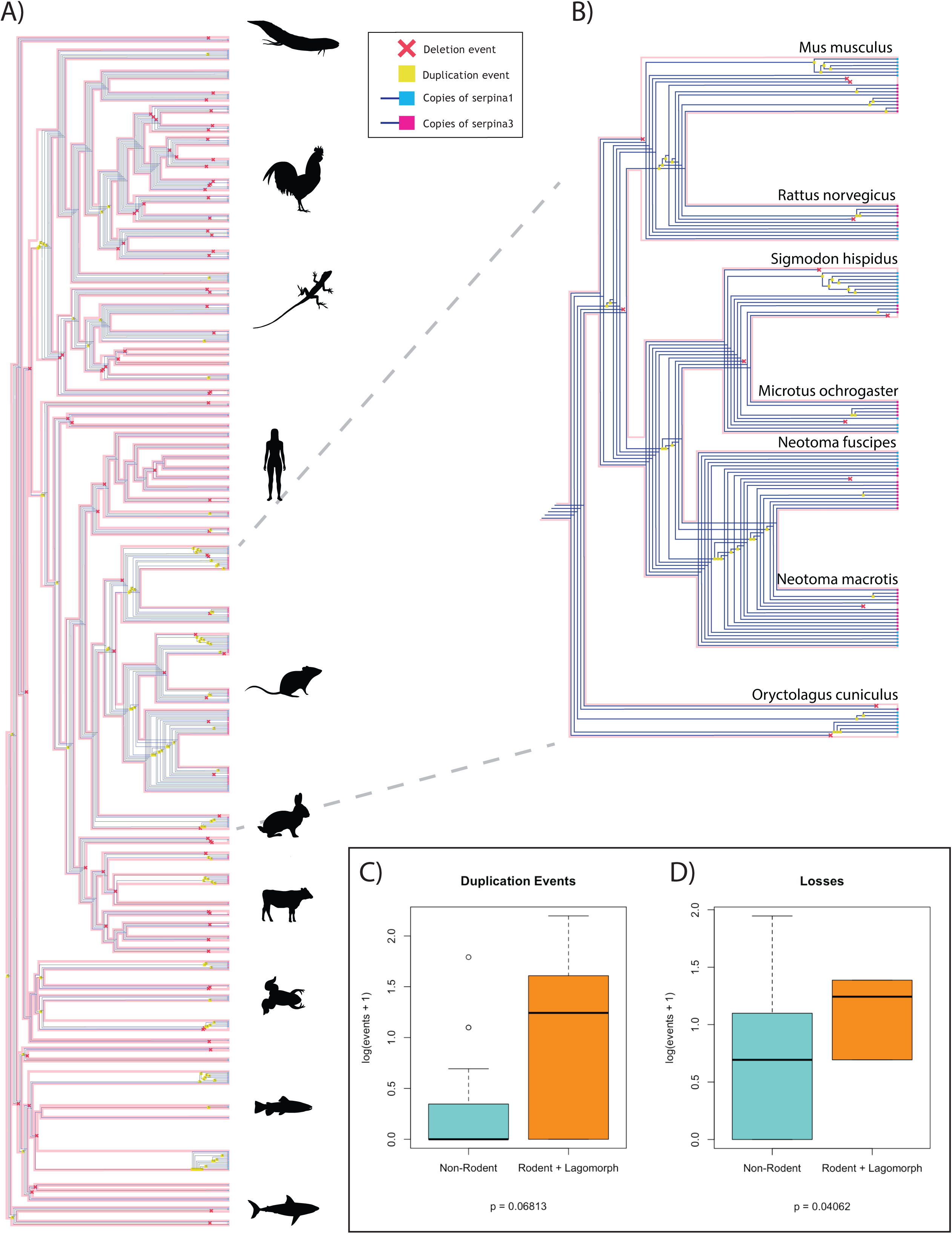
a) Reconciled gene tree created using GeneRax v2.1.3 and visualized using ThirdKind v3.5.0. Duplications are represented by yellow squares, losses by red X’s. The tips of the tree are colored to reflect the identity of the gene (SERPINA1 = cyan; SERPINA3 = pink). Duplications have independently occurred within mammals, fish, reptiles, and birds. b) The rodent clade of the reconciled gene tree. Most duplications occur within genera, many being repeated duplications of the same ancestral gene. c/d) Comparison of SERPIN (C) duplication and (D) loss events in non-rodents vs. rodents and related lagomorphs. Rodents and lagomorphs have more frequent duplications (p=0.068) and losses (0.041) than other species. A log scale was used to account for differences in variance between the two datasets.

We successfully expressed all 12 SERPINA3 paralogs from *N. macrotis* with high purity, while only 2 out of 5 SERPINA1 paralogs were expressed and their purity was low (Fig. S2). We therefore focused largely on SERPINA3 proteins for detailed functional characterization. SERPINA3 paralogs were reacted with cathepsin G, chymotrypsin, and trypsin. If inhibition occurs, we expect a protein band to form that is larger in molecular weight than either the purified serpin or the purified proteases, representing the inhibitory complex. Out of the SERPINA3s, paralogs 3-3, 3-5, 3-8, 3-10, and 3-12 formed complexes with both cathepsin G and chymotrypsin (Fig. 3a, S4). All of these, except SERPINA3-5, are in the same clade, henceforth “clade B;” this clade also includes SERPINA3-7, which did not form complexes with either protease. SERPINA3-5 serves as the outgroup to the remaining SERPINs, henceforth “clade A.” Along with variation in complex formation, SERPINA3 paralogs showed variety in the extent to which they were cleaved at all by the proteases. For example, SERPINA3-6 was almost completely cleaved by chymotrypsin, whereas other SERPINs that do not form complexes are not completely degraded in the same time frame. However, SERPINA3-6 does not appear to have been degraded far beyond the initial cleavage: there are two thick bands slightly below the control band, likely composed of SERPINA3-6 that was cleaved at multiple different sites, then resisted further degradation. In contrast, SERPINA3-1 and SERPINA3-4 were cleaved several times, forming multiple lower molecular weight bands.

**Figure 3.**
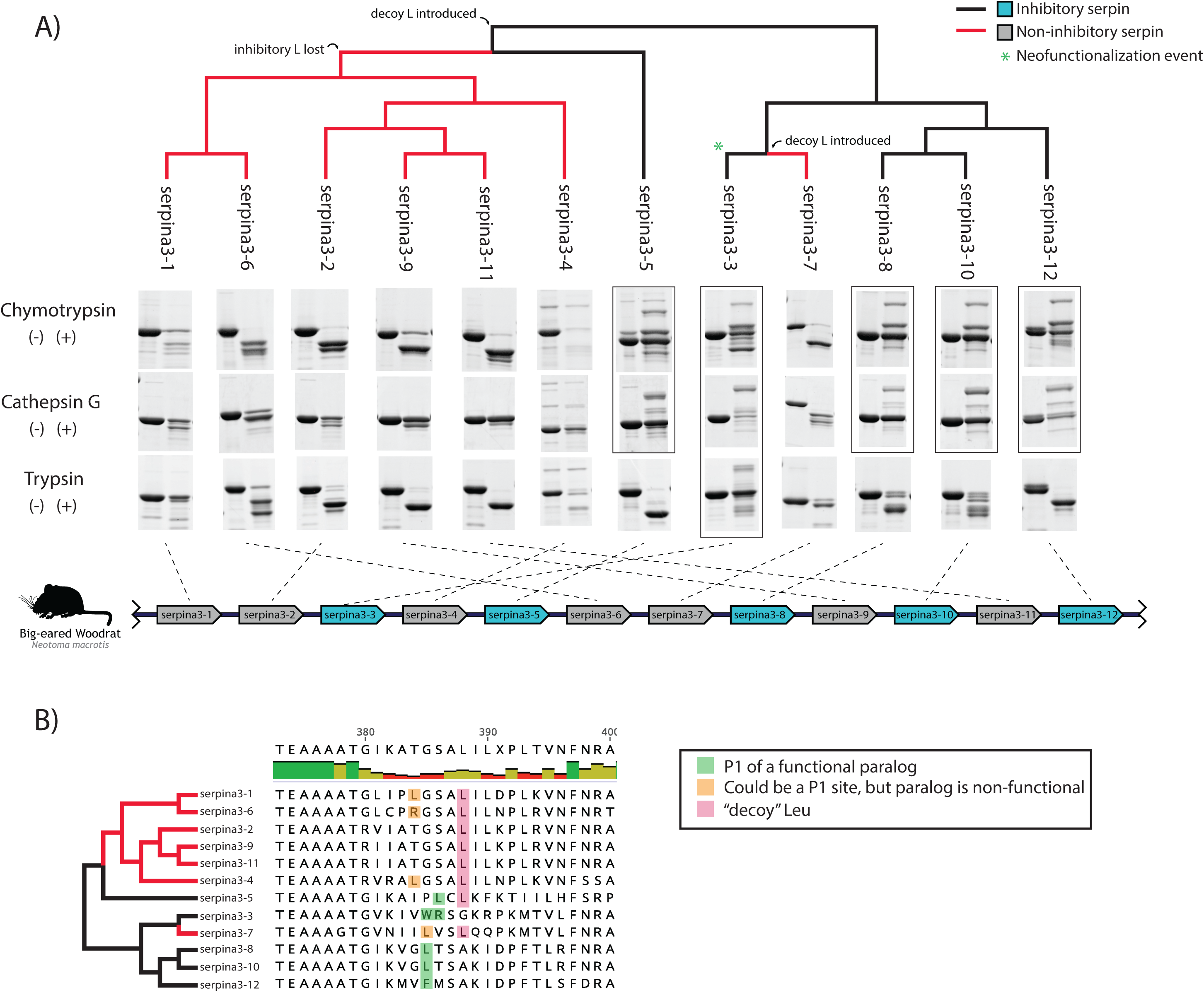
Evolution of SERPINA3-like gene function in *Neotoma macrotis.* (a) Comparison of the function of woodrat SERPINA3 paralogs against chymotrypsin, cathepsin G, and trypsin, mapped onto a phylogeny of gene relationships of woodrat SERPINA3s. Two clades emerged, one that consisted primarily of paralogs that did not inhibit any of the tested targets and one that consisted primarily of inhibitory paralogs. Most of the members of this inhibitory clade only inhibited chymotrypsin and cathepsin G, but SERPINA3-3 underwent a neofunctionalization event to inhibit trypsin. SDS-PAGE images indicating inhibitory complex formation are boxed, and the corresponding genes on the genome are blue. In the left lane of each cropped gel result is the SERPIN-only control, whereas the right lane is the mixture of SERPIN and protease. Protease-only controls and uncropped versions of each result can be found in Figure S4. The genomic orientation of the genes encoding each paralog is indicated by the dotted lines; duplication events do not correspond with synteny on the chromosome. (b) Comparison of the RCL region of all woodrat SERPINA3 paralogs. SERPINA3-3 has consecutive Trp and Arg residues, which we believe is the source of its neofunctionalization to inhibit trypsin as well as cathepsin G and chymotrypsin.

Though the SERPINA3 paralogs that formed complexes with cathepsin G and chymotrypsin were the same, the degradation patterns differed (Fig. 3a, S4). Most paralogs resisted degradation by cathepsin G much more effectively than they resisted degradation by chymotrypsin, despite a longer incubation time with the former. Only SERPINA3-7 was heavily degraded by cathepsin G. This likely reflects the more generalized substrate preferences of chymotrypsin compared to cathepsin G.

One SERPINA3 paralog (3-3) formed an inhibitory complex with trypsin (Fig. 3a, S4). Like other proteases, trypsin shows differential primary and secondary cleavage patterns of SERPINA3 paralogs; most SERPINs were cleaved once but resisted further cleavage. There was no observed difference in cleavage pattern between clades. Relative to the background of chymotrypsin-like protease inhibition in SERPINA3-like genes, trypsin-inhibition of SERPINA3-3 is a significant neofunctionalization event that renders a broad inhibitory capacity in the resulting protein.

We next tested the SERPINA paralogs with the purified RVV-V and Protac. Two paralogs, SERPINA3-3 and SERPINA3-12, formed high molecular weight inhibitory complexes with these SVSPs (Fig. 5). Both SERPINs formed more complex with Protac than with RVV-V, suggesting substrate specificity between individual venom proteins.

Next, SERPINA3 paralogs were reacted with the benzamidine-purified SVSPs from rattlesnake predators of woodrats, Complexes were observed between SERPINA3-3 and isolated SVSPs from C. *oreganus*, *C. molossus*, and *C. adamanteus* (Fig. 4, S5). The most complex was formed between SERPINA3-3 and *C. adamanteus* SVSPs, while the least was formed with the *C. oreganus* SVSPs. Three complexes of different molecular weights appeared in the mixture with *C. molossus*. In addition, a complex was detected between SERPINA3-12 and *C. molossus* SVSPs. All paralogs showed evidence for a substrate pathway, with a band appearing slightly under the molecular weight of the control serpin, suggesting that SVSPs are sometimes escaping following SERPIN cleavage (Fig. S5).

**Figure 4.**
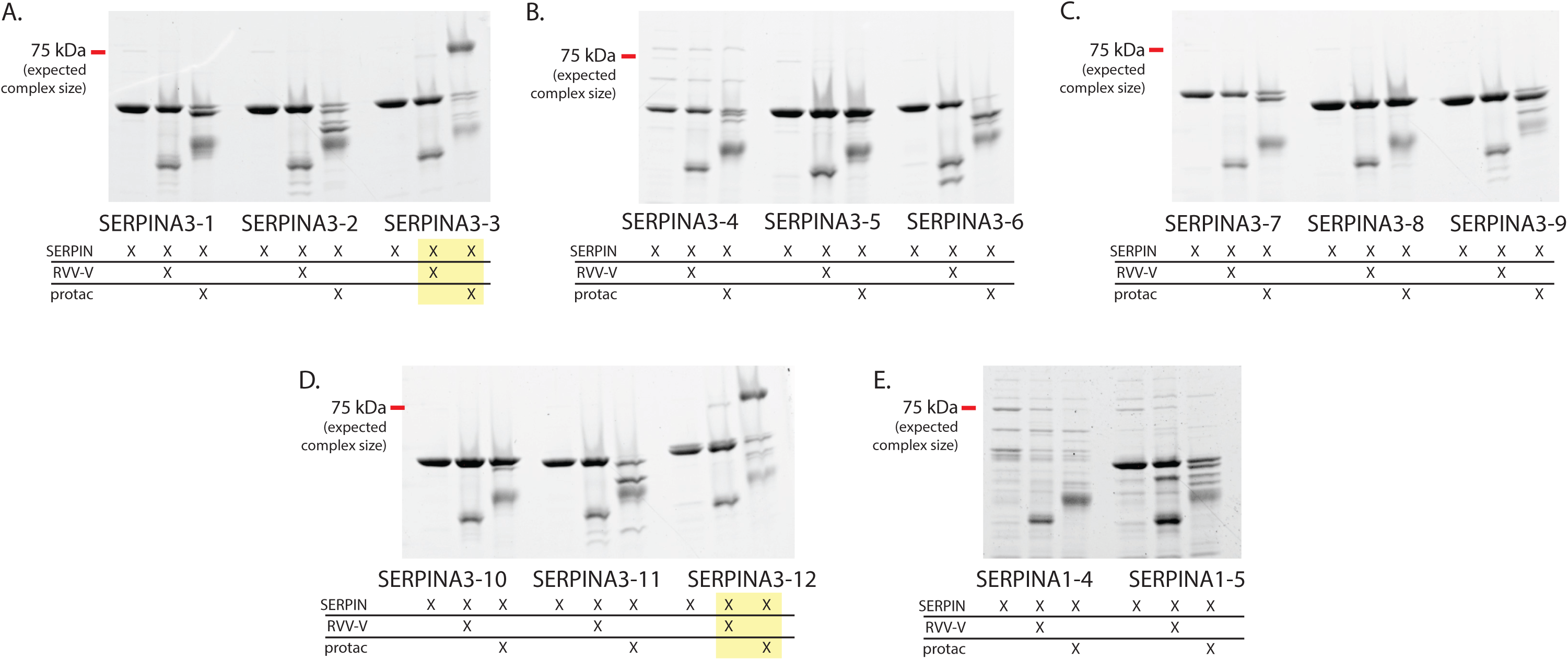
Inhibition of Purified Venom SVSP by SERPINA3-like Proteins: Each of the 12 SERPINA3 paralogs was tested against Russell’s viper venom factor V activator (RVV-V) and *A. contortrix* Protein C Activator (Protac) on Coomassie Blue-stained SDS-PAGE gels. SERPINA3-3 and SERPINA3-12 (highlighted) form complexes with both SVSPs but more readily react with Protac than RVV-V. Uncropped versions of these gels are provided in Fig. S6. The X marks below each lane specify the lane contents as either purified SERPIN or SERPIN + SVSP.

## Discussion

Even in laboratory mice, the specific roles for the multiple paralogs of SERPINA1 and SERPINA3 are poorly characterized (Forsyth et al., 2003). We have demonstrated both uniqueness of rodents regarding rates of birth-death evolution of SERPINA genes, as well as empirically demonstrated repeated functional diversification across these tandem gene arrays in a wild rodent examplar. Our examination of general anti-protease activity from selected, non-rodent proteases limits our inferences regarding the exact physiological roles played by these SERPINs in nature. Yet, we do find support for our working hypothesis that at least some of these duplications serve as venom inhibitors, expanding the known functional repertoire of venom-inhibitory proteins to SERPINA3-like paralogs. Generally, rodent SERPINA3s show repeated gains and losses of chymotrypsin-like inhibitory activity, as well as one neofunctionalization event toward dual inhibition of trypsin and chymotrypsin, both of which may be conducive to coevolution with snake venom proteases which can be trypsin-like or chymotrypsin-like at their active sites (Whittington et al., 2018).

### History of Gene Duplications in the SERPINA Locus

In humans, SERPINA1 is one of the most abundant plasma proteins, where it exists as a single copy gene whose product inhibits elastases (Gettins, 2002). Similarly, SERPINA3 also exists as a single-copy gene in humans and primarily inhibits cathepsin G (Gettins, 2002). Our analysis of gene gains and losses suggest that the large tandem arrays of SERPINA1 and SERPINA3 found in rodents developed via several independent gene duplication events, rather than stemming from one large ancestral duplication event. In addition, there is species-specificity in which gene is most common between SERPINA1 to SERPINA3, with the cotton rat having more SERPINA1s than SERPINA3s, the woodrat and house mouse having more SERPINA3s than A1s, and the prairie vole and Norway rat having similar counts of both. Similar to previous studies in *M. musculus*, the *N. macrotis* SERPINA3 duplications as determined phylogenetically show little correlation with the orientation or order of genes on the chromosome (Inglis and Hill, 1991). However, woodrat SERPINA1 paralogs are more divergent, on average, than the mouse paralogs. Our finding that most duplications occurred within rodent genera is consistent with previous studies that claim that duplications in mice primarily occurred at the shared ancestor of the *Mus* genus (Rheaume et al., 1994).

### Functional diversification

In the SERPINA3s tested in this study, SERPINA3-5, 3-8, and 3-10 have a Leu residue within one residue of the typical P1-P1’ region as observed on a multiple alignment with human SERPINA3 (Fig. 3b). Furthermore, SERPINA3-3 has a Trp residue in the same place, and SERPINA3-12 has a Phe residue. Chymotrypsin cleaves after Trp, Tyr, or Phe, with secondary affinity for cutting after Leu, Met, or His (Keil, 2012). Mapping of complex formation to the of *N. macrotis*’s SERPINA3 gene phylogeny showed a pattern suggesting that ancestral SERPINA3s likely had the ability to form a complex with and inhibit chymotrypsin, and that the ability was lost twice. The first of these two losses to occur was within the clade composed of SERPINA3-1, 3-2, 3-4, 3-5, 3-6, 3-9, and 3-11 (henceforth “clade A”). SERPINA3-5 is the only member of clade A to have retained the ability to inhibit chymotrypsin. Clade A is characterized by the presence of a Leu residue in position 388 on the consensus sequence. Though Leu is a known cleavage site for chymotrypsin, L388 may be too far toward the C-terminal end of the RCL (Fig. S7) to facilitate the trapping action necessary for inhibition of a protease (Jumper et al., 2021). Only SERPINA3-5 was able to form a complex while also having a leucine residue at position 388. However, it also has another Leu residue shortly before L388, which we hypothesize serves as a kinetically favored cleavage target for chymotrypsin. Previous research has shown that certain SERPINA1 paralogs with seemingly valid P1-P1’ sites are unable to form a complex with chymotrypsin, likely due to interference by other residues in the RCL that prevent or hinder the insertion of the RCL into the central β-sheet (Barbour et al., 2002a).

The second loss of chymotrypsin inhibitory function occurred in clade B. Most of the paralogs in this clade retain inhibitory function, but SERPINA3-7 has lost that ability. As stated before, there are several different functional residues at the P1 site in clade B: Trp, Leu, and Phe, all at position 385 in the consensus. SERPINA3-7 also has the L385 residue but has also separately evolved the inhibitory Leu at position 388. This serves as further evidence that L388 can block inhibitory function, though there could be other explanations for this loss of function in SERPINA3-7, like the fact that P1’ in SERPINA3-7 is Val, whereas P1’ in all other functional paralogs is a polar amino acid (Cys, Arg, Thr, or Met).

Cathepsin G cleaves after Trp, Tyr, Phe, and Leu, with secondary action after Lys and low activity after Arg: similar cleavage sites to chymotrypsin (Thorpe et al., 2018). Thus, it is unsurprising that the same paralogs formed complexes with cathepsin G and chymotrypsin. However, the extent to which the SERPINs entered the substrate pathway did differ between the two proteases: SERPINs showed much greater resistance to primary and secondary cleavage by cathepsin G than they did chymotrypsin. This could be tied to cathepsin G’s substrate specificities being stricter than chymotrypsin’s (Polanowska et al., 1998).

Trypsin cleaves after Lys and Arg residues, unless P1’ is Pro (Manea et al., 2007). In human SERPINA1, Met at P1 also successfully inhibits trypsin (Gettins and Olson, 2009; Huntington, 2011). Previous studies in mouse SERPINA1 paralogs have demonstrated even more freedom in the P1 site, with Tyr-Ser also serving as a valid cleavage site (Barbour et al., 2002a). Despite this apparent relative flexibility, in this study, only SERPINA3-3 formed a complex with trypsin, having evolved an Arg at position 386 (Fig. 3b). This indicates a neofunctionalization event where SERPINA3-3 evolved to inhibit trypsin while retaining the ability to inhibit chymotrypsin. However, most of the other SERPINA3 paralogs show no activity against trypsin. The only other paralog that has a potential cleavage site is SERPINA3-4, with an Arg at position 384, though it appears not to form a complex with trypsin. The minimal trypsin target sites among SERPINA3 paralogs in *N. macrotis* is in strong contrast to previous studies into *M. musculus* SERPINA3, which found as many trypsin cleavage sites as there were chymotrypsin cleavage sites, with no overlap (Horvath et al., 2004). As such, it cannot be assumed that rodents with similar patterns of copy number evolution will demonstrate similar protease inhibition profiles.

### Inhibition of Snake Venom Serine Proteases

In the genus *Mus,* (Barbour et al., 2002a) showed that their five SERPINA1 paralogs are capable of SVSP inhibition, with each paralog forming unique patterns of complex depending on the snake species’ venom present. A major implication of our work is to extend these results beyond SERPINA1 paralogs to the SERPINA3 paralogs, which are often more gene copy-rich than SERPINA1. One limitation of our study that could have reduced the SVSP-inhibiting ability of our SERPINs was the lack of glycosylation on our engineered proteins; many natural SERPINs are glycosylated. Certain types of fucosylation have been found to regulate the ability of SERPINA1 to bind to elastase, promoting or inhibiting activity depending on the type of terminal monosaccharide (D. Wu et al., 2023). The *M. saxicola* SERPINA1 paralogs were expressed in mammalian COS-1 cells (Barbour et al., 2002a) rather than *E. coli*, and may therefore have had more natural glycosylation patterns. While our results with non-glycosylated proteins may not reflect in the entirety the SERPINA-based defense mechanisms of *N. macrotis* against snake venom, they still offer a large step in showing that SERPINA3-like proteins are also capable of inhibiting diverse SVSPs.

Most of the snake species sympatric with *N. macrotis* have primarily trypsin-like serine proteases (Whittington et al., 2018), as with all the venoms we tested herein. SERPINA3-3’s neofunctionalization to a trypsin-like inhibitor may thus contribute to natural inhibition of these trypsin-like venoms (Fig. 4). Though SERPINA3-12 lacks a Lys or Arg to serve as a trypsin cleavage site, it also successfully formed a complex with the C. *molossus* SVSPs. (Fig. 3b, 6). However, rates of inhibition appear to be significantly slower than SERPINA3-3, requiring a long exposure to be visualized. This could be due to the relatively small amount of susceptible SVSP isoforms loaded: the SVSP fraction was a mixture of many different serine proteases, and venom metalloproteinases likely were present as contaminants as well. The identity of the exact SVSPs in the venom mixture inhibited by each SERPIN may differ as well and, in the case of SERPINA3-3 and *C. molossus* SVSPs, one SERPIN paralog appears to complex with multiple isoforms of SVSPs.

Woodrat SERPINA3-3 formed complexes with SVSPs in all venoms tested, while the degree of complex formation varied. The most complex was formed with *C. adamanteus* and the least with *C. oreganus*. This pattern is consistent with coevolutionary local adaptation of the snakes to avoiding woodrat inhibitors, since *C. oreganus* is the local predator of these woodrats while *C. adamanteus* is found on the opposite side of the continent. This corroborates previous work in other woodrat-snake systems that suggested that the snake is winning the proverbial “molecular arms race” (Robinson et al., 2021). This conclusion has some limits, as seen in the experiments with RVV-V and Protac (Fig. 5). Russell’s Viper is native to India and Bangladesh while *A. contortrix* is native to Eastern North America (Guiher and Burbrink, 2008; Wüster et al., 1992). The woodrat’s SERPINA3 paralogs were more effective inhibitors of Protac from *A. contortrix* than RVV-V from Russell’s Viper, suggesting potential adaptation to inhibit venoms from the clade of vipers (Western Hemisphere Crotalinae) with which they have coevolved compared to Eastern Hemisphere Viperinae like Russell’s Viper. Such patterns of differential susceptibility and adaptation across phylogenetic scales are consistent with findings for snake venom metalloproteinase inhibition by squirrels (Pomento et al., 2016), as well as patterns of host and non-host resistance in host-pathogen systems (Antonovics et al., 2013)

**Figure 5.**
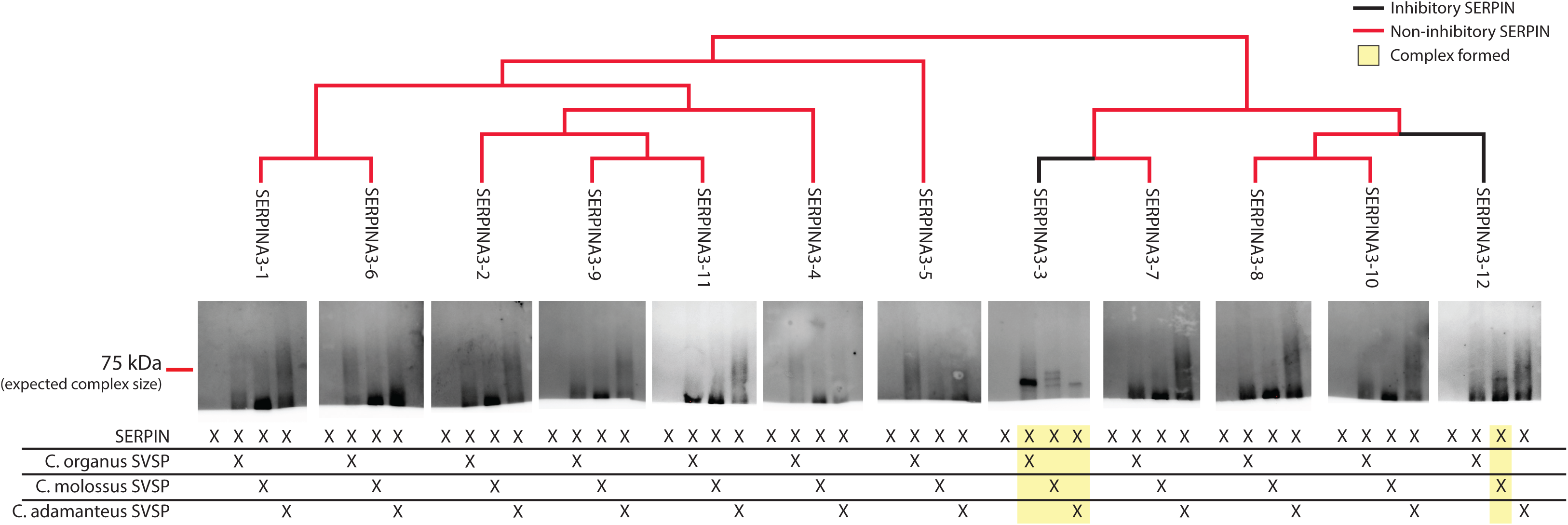
Complex Formation with SVSP from Rattlesnake Predators. These western blots visualize the potential of SERPINA3 paralogs to form inhibitory complexes with isolated SVSPs from three different rattlesnakes, and are ordered according to the phylogenetic relationships of woodrat SERPINA3 paralogs as indicated by the tree. The reaction took place in a 5:2 ratio of SERPIN to SVSP and at 37L. SERPINA3-3 formed a complex with all three snakes’ SVSPs, whereas SERPINA3-12 only formed a complex with *C. molossus* SVSPs. These images depict the portion of each blot above 50 kDa for better exposure of the complexes; full versions of blot images are provided in Figure S5.

In conclusion, the results of this study show how frequent gene duplication can produce functional variation, especially in SERPINs, where the residues of the RCL near the scissile bond can have such a strong effect on the SERPIN’s target. Closest homology to human single-copy orthologs — especially homology of the entire sequence rather than the RCL — is therefore an unreliable indicator of function given the one-to-many gene relationships between species and potential for rapid neofunctionalization via one or a few mutations. In addition, we have provided more evidence that tandem duplication can have a role in complex coevolving systems like venom toxicity and resistance, and employed a systematic testing method applicable to future work in characterizing other tandemly arrayed genes. SERPINA3 paralogs should be added to the growing list of resistance factors that protect mammals in molecular warfare with venomous snakes.

## Acknowledgements

This research was supported by the University of Michigan’s Honors Summer Fellowship and NIH grant 1R35-HL171421. We are also indebted to Marjorie Matocq for supplying the unpublished genomes from *N. fuscipes* and *N. macrotis*, as well as Mark Margres for the unpublished genome of *S. hispidus*. We thank members of the Ginsburg Lab for advice and instruction throughout the research process.

## References

1. Almagro Armenteros, J.J., Tsirigos, K.D., Sønderby, C.K., Petersen, T.N., Winther, O., Brunak, S., von Heijne, G., Nielsen, H., 2019. SignalP 5.0 improves signal peptide predictions using deep neural networks. Nature biotechnology 37, 420–423.

2. Altschul, S.F., Gish, W., Miller, W., Myers, E.W., Lipman, D.J., 1990. Basic local alignment search tool. Journal of Molecular Biology 215, 403–410. 10.1016/S0022-2836(05)80360-2

3. Antonovics, J., Boots, M., Ebert, D., Koskella, B., Poss, M., Sadd, B.M., 2013. The Origin of Specificity by Means of Natural Selection: Evolved and Nonhost Resistance in Host– Pathogen Interactions. Evolution 67, 1–9. 10.1111/j.1558-5646.2012.01793.x

4. Balchan, N.R., Smith, C.F., Mackessy, S.P., 2024. A plethora of rodents: Rattlesnake predators generate unanticipated patterns of venom resistance in a grassland ecosystem. Toxicon: X 21, 100179. 10.1016/j.toxcx.2023.100179

5. Barbour, K.W., Goodwin, R.L., Guillonneau, F., Wang, Y., Baumann, H., Berger, F.G., 2002a. Functional Diversification During Evolution of the Murine α1-Proteinase Inhibitor Family: Role of the Hypervariable Reactive Center Loop. Molecular Biology and Evolution 19, 718–727. 10.1093/oxfordjournals.molbev.a004130

6. Barbour, K.W., Wei, F., Brannan, C., Flotte, T.R., Baumann, H., Berger, F.G., 2002b. The Murine α1-Proteinase Inhibitor Gene Family: Polymorphism, Chromosomal Location, and Structure. Genomics 80, 515–522. 10.1006/geno.2002.6864

7. Benkman, C.W., Parchman, T.L., Favis, A., Siepielski, A.M., 2003. Reciprocal selection causes a coevolutionary arms race between crossbills and lodgepole pine. Am Nat 162, 182–194. 10.1086/376580

8. Biardi, J.E., Coss, R.G., Smith, D.G., 2000. California ground squirrel (*Spermophilus beecheyi*) blood sera inhibits crotalid venom proteolytic activity. Toxicon 38, 713–721. 10.1016/S0041-0101(99)00179-8

9. Borriello, F., Krauter, K.S., 1991. Multiple murine alpha 1-protease inhibitor genes show unusual evolutionary divergence. Proceedings of the National Academy of Sciences 88, 9417– 9421. 10.1073/pnas.88.21.9417

10. Brodie, Edmund D., Ridenhour, B.J., Brodie, E. D., 2002. The evolutionary response of predators to dangerous prey: hotspots and coldspots in the geographic mosaic of coevolution between garter snakes and newts. Evolution 56, 2067–2082. 10.1111/j.0014-3820.2002.tb00132.x

11. Brovkina, M.V., Chapman, M.A., Holding, M.L., Clowney, E.J., 2023. Emergence and influence of sequence bias in evolutionarily malleable, mammalian tandem arrays. BMC Biol 21, 179. 10.1186/s12915-023-01673-4

12. Casewell, N.R., Wüster, W., Vonk, F.J., Harrison, R.A., Fry, B.G., 2013. Complex cocktails: the evolutionary novelty of venoms. Trends in Ecology & Evolution 28, 219–229. 10.1016/j.tree.2012.10.020

13. Cavalcante, J.D.S., de Almeida, C.A.S., Clasen, M.A., da Silva, E.L., de Barros, L.C., Marinho, A.D., Rossini, B.C., Marino, C.L., Carvalho, P.C., Jorge, R.J.B., Dos Santos, L.D., 2022. A fingerprint of plasma proteome alteration after local tissue damage induced by Bothrops leucurus snake venom in mice. J Proteomics 253, 104464. 10.1016/j.jprot.2021.104464

14. de Wit, C.A., 1982. Resistance of the prairie vole (*Microtus ochrogaster*) and the woodrat (*Neotoma floridana*), in Kansas, to venom of the Osage copperhead (*Agkistrodon contortrix phaeogaster*). Toxicon 20, 709–714. 10.1016/0041-0101(82)90119-2

15. de Wit, C.A., Weström, B.R., 1987. Venom resistance in the Hedgehog, *Erinaceus europaeus*: Purification and identification of macroglobulin inhibitors as plasma antihemorrhagic factors. Toxicon 25, 315–323. 10.1016/0041-0101(87)90260-1

16. Demuth, J.P., Hahn, M.W., 2009. The life and death of gene families. BioEssays 31, 29–39. 10.1002/bies.080085

17. Dowell, N.L., Giorgianni, M.W., Kassner, V.A., Selegue, J.E., Sanchez, E.E., Carroll, S.B., 2016. The Deep Origin and Recent Loss of Venom Toxin Genes in Rattlesnakes. Current Biology 26, 2434–2445. 10.1016/j.cub.2016.07.038

18. Edgar, R.C., 2004. MUSCLE: multiple sequence alignment with high accuracy and high throughput. Nucleic Acids Res 32, 1792–1797. 10.1093/nar/gkh340

19. Forsyth, S., Horvath, A., Coughlin, P., 2003. A review and comparison of the murine α1-antitrypsin and α1-antichymotrypsin multigene clusters with the human clade A serpins. Genomics 81, 336–345. 10.1016/S0888-7543(02)00041-1

20. Gettins, P.G.W., 2002. Serpin Structure, Mechanism, and Function. Chem. Rev. 102, 4751– 4804. 10.1021/cr010170+

21. Gettins, P.G.W., Olson, S.T., 2009. Exosite Determinants of Serpin Specificity. J Biol Chem 284, 20441–20445. 10.1074/jbc.R800064200

22. Gibbs, H.L., Sanz, L., Pérez, A., Ochoa, A., Hassinger, A.T.B., Holding, M.L., Calvete, J.J., 2020. The molecular basis of venom resistance in a rattlesnake-squirrel predator-prey system. Molecular Ecology 29, 2871–2888. 10.1111/mec.15529

23. Guiher, T.J., Burbrink, F.T., 2008. Demographic and phylogeographic histories of two venomous North American snakes of the genus *Agkistrodon*. Molecular Phylogenetics and Evolution 48, 543–553. 10.1016/j.ympev.2008.04.008

24. Hahn, M.W., 2009. Distinguishing Among Evolutionary Models for the Maintenance of Gene Duplicates. J Hered 100, 605–617. 10.1093/jhered/esp047

25. Heit, C., Jackson, B.C., McAndrews, M., Wright, M.W., Thompson, D.C., Silverman, G.A., Nebert, D.W., Vasiliou, V., 2013. Update of the human and mouse SERPINgene superfamily. Human Genomics 7, 22. 10.1186/1479-7364-7-22

26. Holding, M.L., Biardi, J.E., Gibbs, H.L., 2016a. Coevolution of venom function and venom resistance in a rattlesnake predator and its squirrel prey. Proceedings of the Royal Society B: Biological Sciences 283, 20152841. 10.1098/rspb.2015.2841

27. Holding, M.L., Drabeck, D.H., Jansa, S.A., Gibbs, H.L., 2016b. Venom Resistance as a Model for Understanding the Molecular Basis of Complex Coevolutionary Adaptations. Integrative and Comparative Biology 56, 1032–1043. 10.1093/icb/icw082

28. Horvath, A.J., Forsyth, S.L., Coughlin, P.B., 2004. Expression Patterns of Murine Antichymotrypsin-like Genes Reflect Evolutionary Divergence at the Serpina3 Locus. J Mol Evol 59, 488–497. 10.1007/s00239-004-2640-9

29. Huntington, J.A., 2011. Serpin structure, function and dysfunction. Journal of Thrombosis and Haemostasis 9, 26–34. 10.1111/j.1538-7836.2011.04360.x

30. Inglis, J.D., Hill, R.E., 1991. The murine Spi-2 proteinase inhibitor locus: a multigene family with a hypervariable reactive site domain. EMBO J 10, 255–261. 10.1002/j.1460-2075.1991.tb07945.x

31. Itoh, N., Tanaka, N., Funakoshi, I., Kawasaki, T., Mihashi, S., Yamashina, I., 1988. Organization of the gene for batroxobin, a thrombin-like snake venom enzyme. Homology with the trypsin/kallikrein gene family. Journal of Biological Chemistry 263, 7628–7631. 10.1016/S0021-9258(18)68544-8

32. Jumper, J., Evans, R., Pritzel, A., Green, T., Figurnov, M., Ronneberger, O., Tunyasuvunakool, K., Bates, R., Žídek, A., Potapenko, A., Bridgland, A., Meyer, C., Kohl, S.A.A., Ballard, A.J., Cowie, A., Romera-Paredes, B., Nikolov, S., Jain, R., Adler, J., Back, T., Petersen, S., Reiman, D., Clancy, E., Zielinski, M., Steinegger, M., Pacholska, M., Berghammer, T., Bodenstein, S., Silver, D., Vinyals, O., Senior, A.W., Kavukcuoglu, K., Kohli, P., Hassabis, D., 2021. Highly accurate protein structure prediction with AlphaFold. Nature 596, 583–589. 10.1038/s41586-021-03819-2

33. Keil, B., 2012. Specificity of Proteolysis. Springer Science & Business Media.

34. Kim, D., Paggi, J.M., Park, C., Bennett, C., Salzberg, S.L., 2019. Graph-based genome alignment and genotyping with HISAT2 and HISAT-genotype. Nat Biotechnol 37, 907– 915. 10.1038/s41587-019-0201-4

35. Kumar, S., Suleski, M., Craig, J.M., Kasprowicz, A.E., Sanderford, M., Li, M., Stecher, G., Hedges, S.B., 2022. TimeTree 5: An Expanded Resource for Species Divergence Times. Molecular Biology and Evolution 39, msac174. 10.1093/molbev/msac174

36. Lallemand, T., Leduc, M., Landès, C., Rizzon, C., Lerat, E., 2020. An Overview of Duplicated Gene Detection Methods: Why the Duplication Mechanism Has to Be Accounted for in Their Choice. Genes (Basel) 11, 1046. 10.3390/genes11091046

37. Lawrence, D.A., Ginsburg, D., Day, D.E., Berkenpas, M.B., Verhamme, I.M., Kvassman, J.-O., Shore, J.D., 1995. Serpin-Protease Complexes Are Trapped as Stable Acyl-Enzyme Intermediates (∗). Journal of Biological Chemistry 270, 25309–25312. 10.1074/jbc.270.43.25309

38. Markland, F.S., 1998. Snake venoms and the hemostatic system. Toxicon 36, 1749–1800. 10.1016/S0041-0101(98)00126-3

39. Morel, B., Kozlov, A.M., Stamatakis, A., Szöllősi, G.J., 2020. GeneRax: A Tool for Species-Tree-Aware Maximum Likelihood-Based Gene Family Tree Inference under Gene Duplication, Transfer, and Loss. Mol Biol Evol 37, 2763–2774. 10.1093/molbev/msaa141

40. Ochoa, A., Hassinger, A.T.B., Holding, M.L., Gibbs, H.L., 2023. Genetic characterization of potential venom resistance proteins in California ground squirrels (Otospermophilus beecheyi) using transcriptome analyses. Journal of Experimental Zoology Part B: Molecular and Developmental Evolution 340, 259–269. 10.1002/jez.b.23145

41. Omori-Satoh, T., Yamakawa, Y., Mebs, D., 2000. The antihemorrhagic factor, erinacin, from the European hedgehog (*Erinaceus europaeus*), a metalloprotease inhibitor of large molecular size possessing ficolin/opsonin P35 lectin domains. Toxicon 38, 1561–1580. 10.1016/S0041-0101(00)00090-8

42. Penel, S., Menet, H., Tricou, T., Daubin, V., Tannier, E., 2022. Thirdkind: displaying phylogenetic encounters beyond 2-level reconciliation. Bioinformatics 38, 2350–2352. 10.1093/bioinformatics/btac062

43. Perez, J.C., Haws, W.C., Hatch, C.H., 1978. Resistance of woodrats (*Neotoma micropus*) to *Crotalus atrox* venom. Toxicon 16, 198–200. 10.1016/0041-0101(78)90039-9

44. Polanowska, J., Krokoszynska, I., Czapinska, H., Watorek, W., Dadlez, M., Otlewski, J., 1998. Specificity of human cathepsin G. Biochimica et Biophysica Acta (BBA) - Protein Structure and Molecular Enzymology 1386, 189–198. 10.1016/S0167-4838(98)00085-5

45. Pomento, A.M., Perry, B.W., Denton, R.D., Gibbs, H.L., Holding, M.L., 2016. No safety in the trees: Local and species-level adaptation of an arboreal squirrel to the venom of sympatric rattlesnakes. Toxicon 118, 149–155. 10.1016/j.toxicon.2016.05.003

46. Rheaume, C., Goodwin, R.L., Latimer, J.J., Baumann, H., Bergen, F.G., 1994. Evolution of murine α1-proteinase inhibitors: Gene amplification and reactive center divergence. J Mol Evol 38, 121–131. 10.1007/BF00166159

47. Robinson, K.E., Holding, M.L., Whitford, M.D., Saviola, A.J., Yates, J.R., III, Clark, R.W., 2021. Phenotypic and functional variation in venom and venom resistance of two sympatric rattlesnakes and their prey. Journal of Evolutionary Biology 34, 1447–1465. 10.1111/jeb.13907

48. Sayers, E.W., Bolton, E.E., Brister, J.R., Canese, K., Chan, J., Comeau, D.C., Connor, R., Funk, K., Kelly, C., Kim, S., Madej, T., Marchler-Bauer, A., Lanczycki, C., Lathrop, S., Lu, Z., Thibaud-Nissen, F., Murphy, T., Phan, L., Skripchenko, Y., Tse, T., Wang, J., Williams, R., Trawick, B.W., Pruitt, K.D., Sherry, S.T., 2022. Database resources of the national center for biotechnology information. Nucleic Acids Res 50, D20–D26. 10.1093/nar/gkab1112

49. Scott, B.M., Sheffield, W.P., 2020. Engineering the serpin α1-antitrypsin: A diversity of goals and techniques. Protein Sci 29, 856–871. 10.1002/pro.3794

50. Serrano, S.M.T., 2013. The long road of research on snake venom serine proteinases. Toxicon, Milestones and future prospects in snake venom research 62, 19–26. 10.1016/j.toxicon.2012.09.003

51. The UniProt Consortium, 2023. UniProt: the Universal Protein Knowledgebase in 2023. Nucleic Acids Research 51, D523–D531. 10.1093/nar/gkac1052

52. Thompson, J.D., Higgins, D.G., Gibson, T.J., 1994. CLUSTAL W: improving the sensitivity of progressive multiple sequence alignment through sequence weighting, position-specific gap penalties and weight matrix choice. Nucleic Acids Research 22, 4673–4680. 10.1093/nar/22.22.4673

53. Thompson, J.N., 2005. Coevolution: The Geographic Mosaic of Coevolutionary Arms Races. Current Biology 15, R992–R994. 10.1016/j.cub.2005.11.046

54. Thorpe, M., Fu, Z., Chahal, G., Akula, S., Kervinen, J., de Garavilla, L., Hellman, L., 2018. Extended cleavage specificity of human neutrophil cathepsin G: A low activity protease with dual chymase and tryptase-type specificities. PLoS One 13, e0195077. 10.1371/journal.pone.0195077

55. Toju, H., 2008. Fine-scale local adaptation of weevil mouthpart length and camellia pericarp thickness: altitudinal gradient of a putative arms race. Evolution 62, 1086–1102. 10.1111/j.1558-5646.2008.00341.x

56. Whittington, A.C., Mason, A.J., Rokyta, D.R., 2018. A Single Mutation Unlocks Cascading Exaptations in the Origin of a Potent Pitviper Neurotoxin. Molecular Biology and Evolution 35, 887–898. 10.1093/molbev/msx334

57. Wu, D., Guo, M., Robinson, C.V., 2023. Connecting single-nucleotide polymorphisms, glycosylation status, and interactions of plasma serine protease inhibitors. Chem 9, 665– 681. 10.1016/j.chempr.2022.11.018

58. Wu, Z., Yuan, R., Gu, Q., Wu, X., Gu, L., Ye, X., Zhou, Y., Huang, J., Wang, Z., Chen, X., 2023. Parasitoid Serpins Evolve Novel Functions to Manipulate Host Homeostasis. Molecular Biology and Evolution 40, msad269. 10.1093/molbev/msad269

59. Wüster, W., Otsuka, S., Malhotra, A., Thorpe, R.S., 1992. Population systematics of Russell’s viper: a multivariate study. Biological Journal of the Linnean Society 47, 97–113. 10.1111/j.1095-8312.1992.tb00658.x

