## Supplemental Information for "Tandem duplication of serpin genes yields functional variation and snake venom inhibitors"

### Supplemental Figures

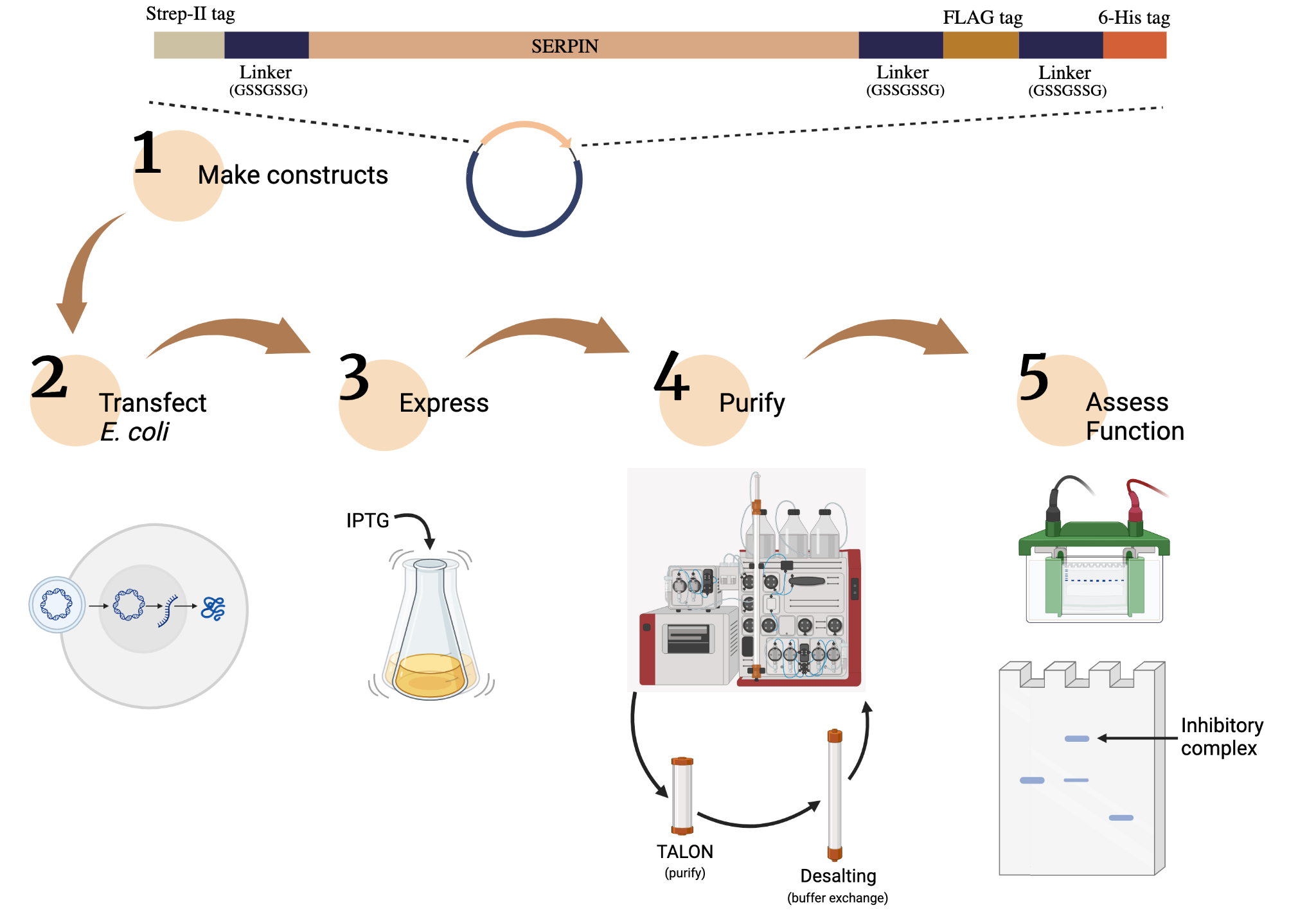

**Figure S1.** Overview of our bacterial expression, purification, and functional study of SERPINA constructs. Constructs were created from all paralogs of SERPINA1 (n=5) and SERPINA3 (n=12) in *N. macrotis*, the big-eared woodrat. Created using BioRender.com.

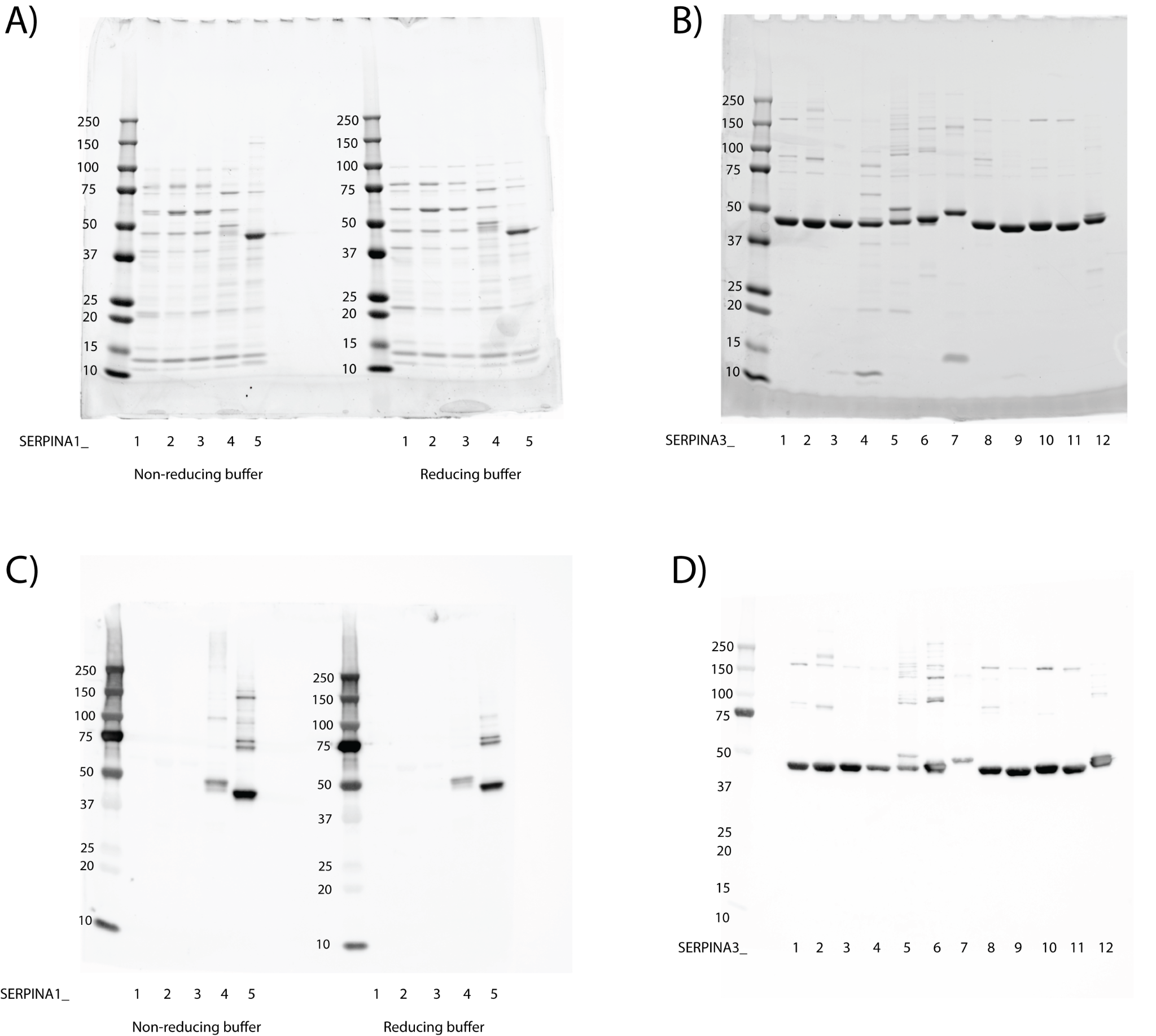

**Figure S2.** Purity analysis of expressed SERPINA1s (a, c) and SERPINA3s (b, d) of *N. macrotis*. Coomassie gels showed low purity in SERPINA1 paralogs (a) but high purity for SERPINA3 paralogs (b). The low purity of SERPINA1 paralogs cannot be attributed to disulfide bonding between SERPINs, as the non-reducing and reducing lanes look almost identical. SERPINA3s were only run with a non-reducing buffer, and most of the impurities appear to be disulfide-bonded SERPINs. Western blots using the N-terminal Strep-II tag showed that SERPINs A1-1, 1-2, and 1-3, were not successfully expressed, and only paralogs 1-4 and 1-5 were present (c). Meanwhile, all SERPINA3s were successfully expressed (d).

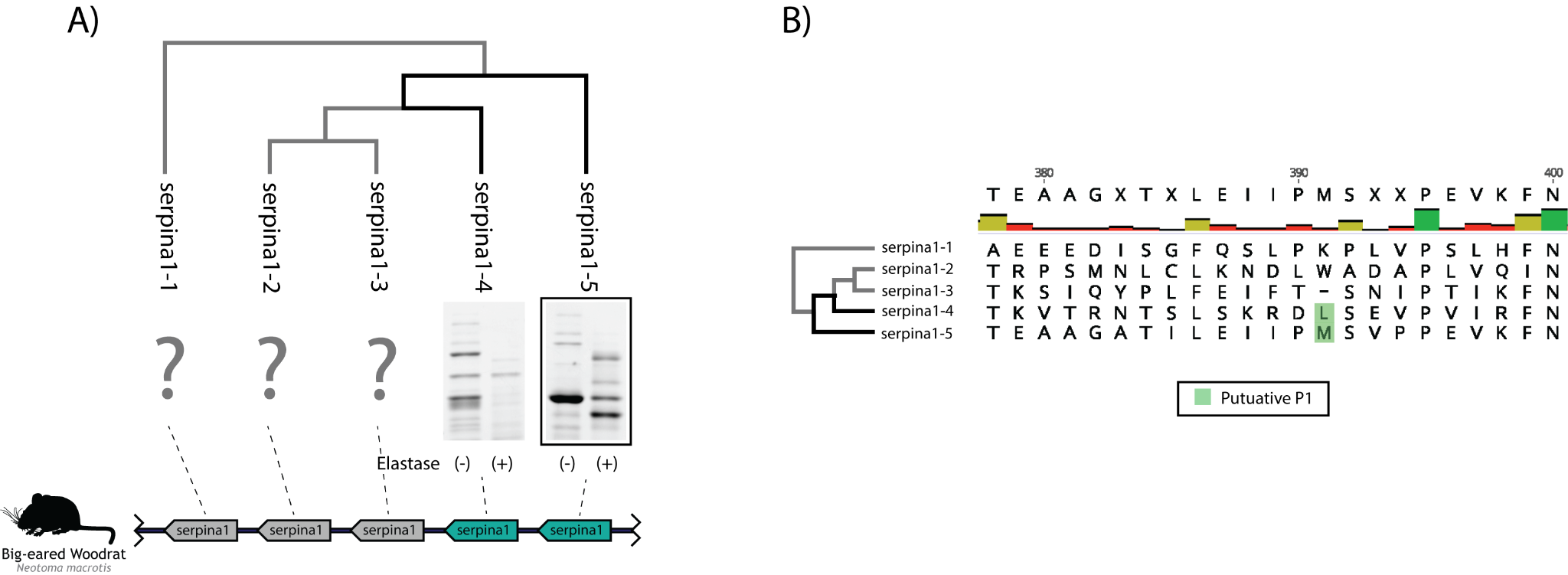

**Figure S3.** (a) Comparison of the function of SERPINA1-4 and 1-5 against elastase, mapped onto a phylogenetic tree of woodrat SERPINA1s. In the left lane of each cropped gel result is the SERPIN-only control, whereas the right lane is the mixture of SERPIN and protease. Protease-only controls and uncropped versions of each result can be found in Figure S4. Both paralogs formed a complex with the elastase at approximately the same molecular weight, with SERPINA1-5’s complex being much darker. The location of the genes encoding each paralog is indicated by the dotted lines; all duplication events resulted in adjacent genes. (b) Comparison of the RCL region of all woodrat SERPINA1 paralogs. Both paralogs’ P1 residue is traditionally capable of facilitating cleavage by elastase, with SERPINA1-5’s M-S P1-P1’ matching that of human SERPINA1. Evolutionary contextualization of SERPINA1 function is difficult without evidence for or against inhibitory ability for the majority of the SERPINA1 paralogs. Upon examining the RCLs of the unrepresented paralogs, few putative cleavage sites for elastase, trypsin, chymotrypsin, or cathepsin G were found. Future work will characterize the function of the unrepresented paralogs and test them as SVSP inhibitors.

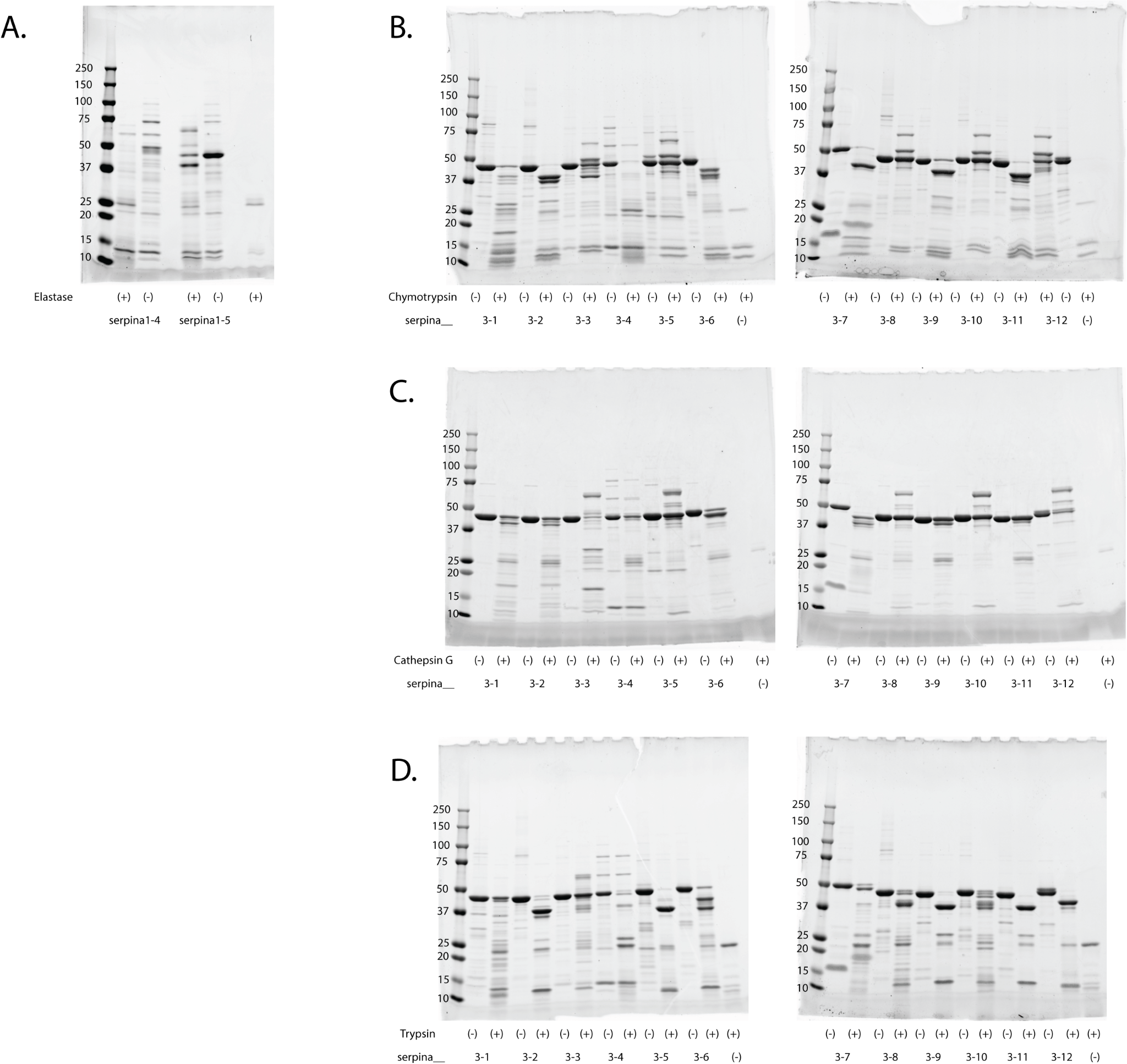

**Figure S4.** Uncropped versions of the Coomassie-stained gels (Fig. 3a) for 2:5 mixtures of neutrophil elastase (a), chymotrypsin (b), cathepsin G (c), and trypsin (d) with SERPINA paralogs.

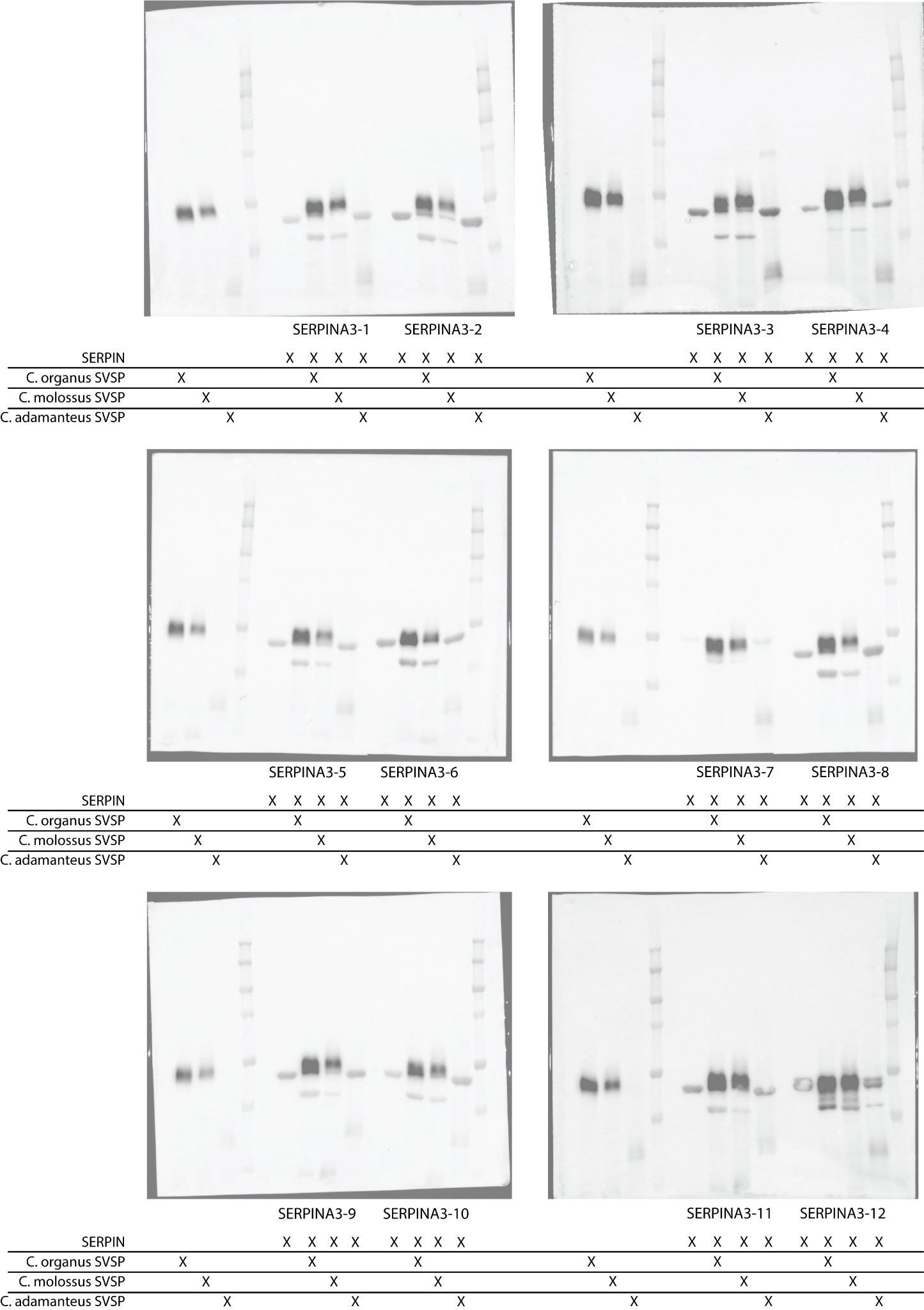

**Figure S5.** Uncropped versions of SERPIN/benzamidine-purified SVSP reaction blots from Figure 5.

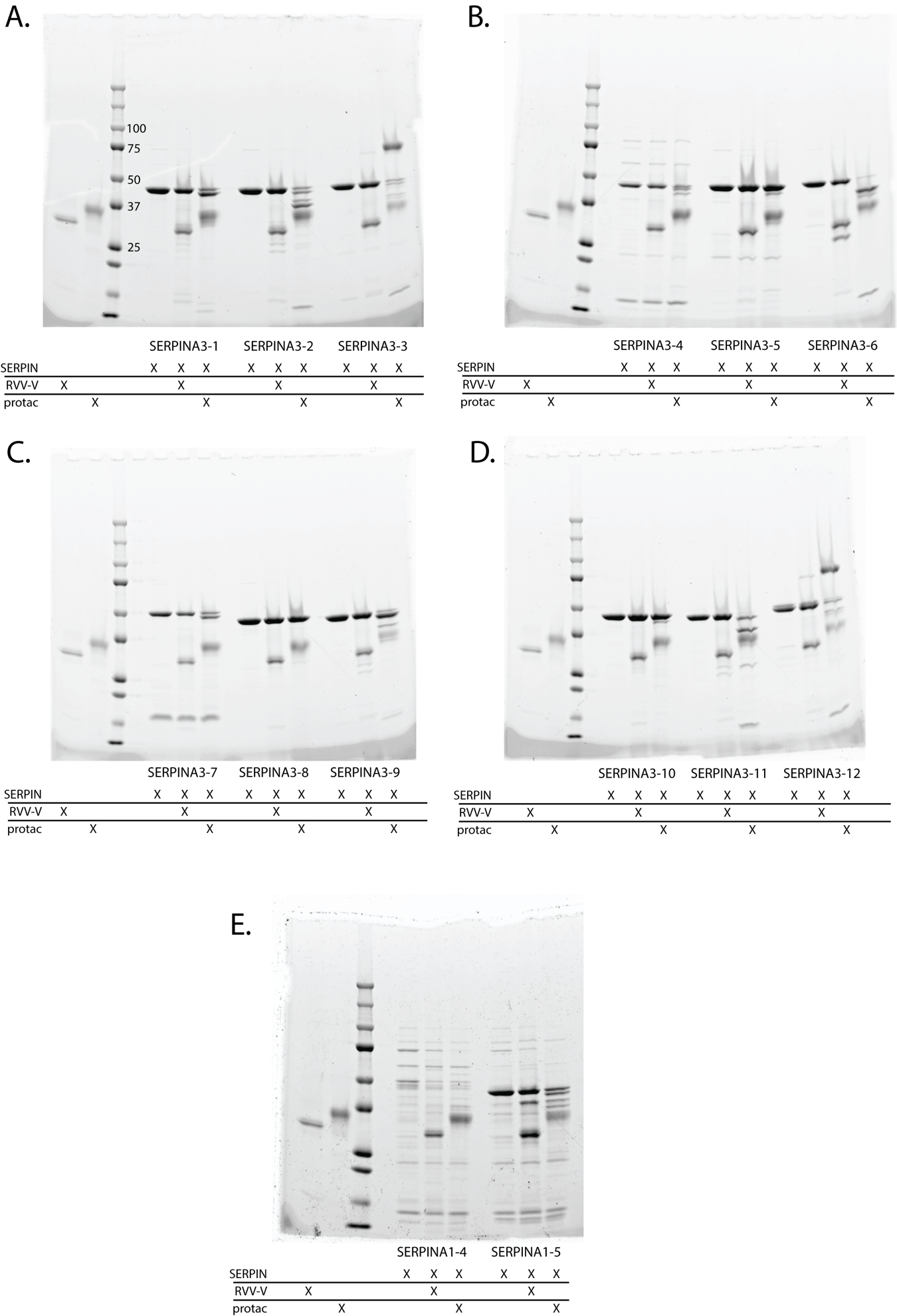

**Figure S6.** Uncropped versions of SERPIN/commercially-purified SVSP reaction blots from Figure 4.

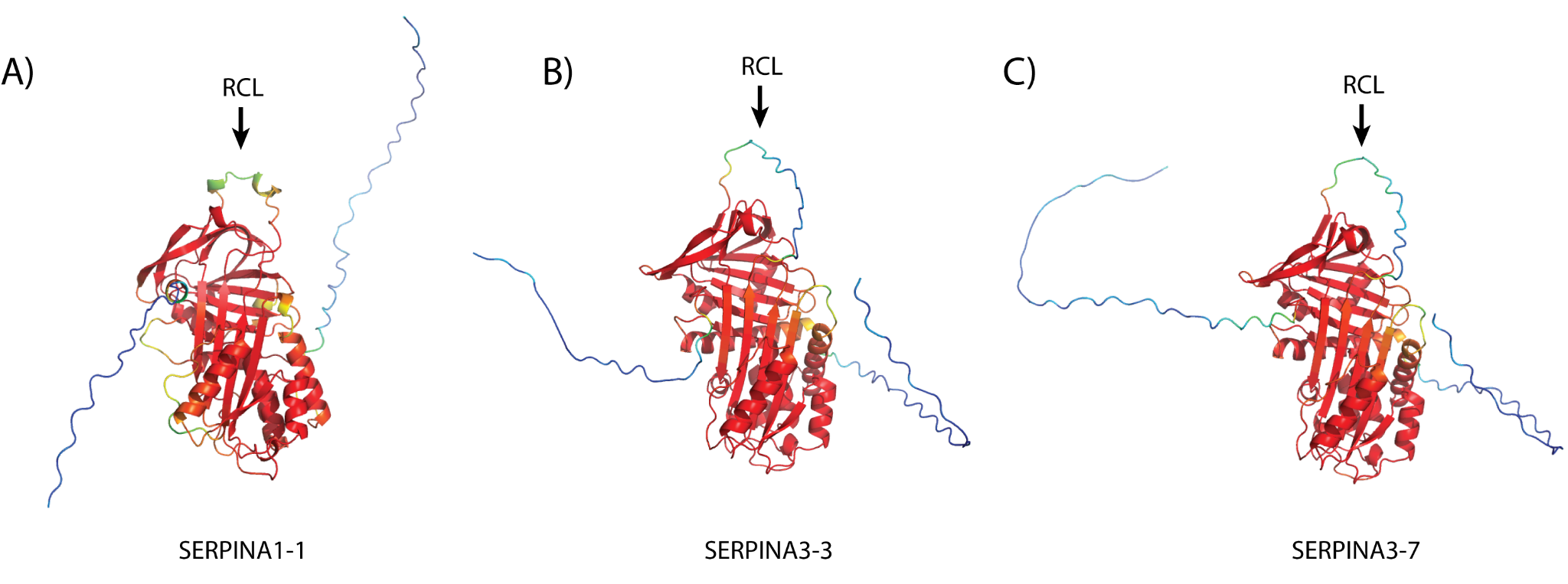

**Figure S7.** Predicted structures of 3 SERPINA paralogs of interest: (a) SERPINA1-1, which was not successfully expressed but doesn’t seem to be a functional inhibitory SERPIN based on the alpha-helix predicted in the RCL; (b) SERPINA3-3, which neofunctionalized to inhibit trypsin as well as chymotrypsin and cathepsin G; and (c) SERPINA3-7, SERPINA3-3’s closest relative in the woodrat, which lost inhibitory function against all proteases tested in this study. Note that these structures include the N-terminal and C-terminal tags, which are not present in the natural sequence of these SERPINs and thus appear as low-confidence predictions on either side of the protein. SERPINA1-1, a more distant relative to other mammal SERPINA1s, contains a Lys residue at position 391. However, this Lys is flanked by two Pro residues, which is an extremely atypical pattern for an RCL; upon simulating the structure of SERPINA1-1, an alpha-helix was predicted in the RCL, which could render that paralog non-functional as an inhibitor through the traditional SERPIN mechanism.
